# Discrimination of conspecific head orientations and learning set formation in Bengalese finches

**DOI:** 10.1101/2022.05.31.494167

**Authors:** Wei Chen, Makiko Kamijo, Kazuo okanoya

**Affiliations:** Department of Life Sciences, Graduate School of Arts and Sciences, the University of Tokyo, Japan; College of Human and Social Science, Kanazawa University, Ishikawa, Japan; Advanced Comprehensive Research Organization, Teikyo University, Tokyo, Japan

**Keywords:** Gaze recognition, generalization, learning set formation

## Abstract

Animals detect others’ gaze direction, such as predators, prey, and courtship mates, to enable adaptive behavior. However, head movements play a more significant role in gaze shifts in birds than eye movement because of the flat eye shape. Additionally, most birds with laterally placed eyes usually turn the lateral face to the target. Therefore, we used Bengalese finches to examine whether songbirds can discriminate between conspecific frontal and lateral faces and divide head orientations in different angles into two categories. In experiments, birds were trained by operant conditioning to discriminate stimuli consisting of conspecific frontal and lateral head images. Eight out of nine birds finished discrimination training and test sessions. The results indicate that Bengalese finches can discriminate gaze direction by the head orientations. Furthermore, as the number of stimuli set increased, fewer trials were needed to achieve the criteria, which suggests that birds have developed a learning set during the training. However, we did not find a clear discrimination boundary for head orientation categorization.

## Introduction

Gaze is an essential cue for studying the mental states of animals, which contains valuable information that reflects the cognitive process, such as selective attention (Dukas, 2002; Yorzinski et al., 2013), causality understanding (Santos & Hauser, 1999), and shared intentions (Tomasello & Carpenter, 2007). Diurnal animals direct the gaze to objects they are paying attention to, including prey, enemies, or courtship partners; on the other hand, they also use the gaze direction as a vital cue to determine whether they are receiving others’ attention so that they can act adaptively. Thus, gaze direction is an ecologically important signal for animals. In recent years, the rapid development of technology to record animal gaze direction has facilitated a vast of studies on eye movement patterns in laboratory environments and when animals are freely behaving. (Haque & Dickman, 2005; Wohlschläger et al., 1993; Pratt, 1982; Yorzinski et al., 2015). However, to understand animal gaze interaction, it is necessary to thoroughly examine the sender’s eye movement patterns and how the receivers perceive the gaze of other individuals.

The utilization of gaze is closely related to the eye’s anatomical structure and visual perception. Like diurnal primates, many diurnal birds also exhibit a high-degree visually dependent behavior. Both primates and birds have high-resolution spots on the retina (a.k.a. the fovea; Hughes, 1977; Walls, 1942) where photoreceptor cells are densely localized. By directing gaze to the object of interest, animals can use their sensitive spots to observe the targets meticulously. Therefore, birds and primates frequently shift their gaze when looking for something.

The base structure of the avian eye is similar to many primates except three main differences: the position of eyes, the structure of the retina, and the eye movements within the orbit. Most primates have forward-facing eyes with only one pair of the fovea, and their gaze is determined only by the projection of the pupils’ centers. Moreover, primates’ eyeballs are structurally round and thus able to move in a wide range. In contrast, for most birds, the eyeballs are located on the sides of the head, and the structure varies in species. Furthermore, those birds have two or more fovea in the retina (raptors, pigeons, and crows have two, most seabirds have one or two, and most sparrows have only one in each retina), and their eyeballs are large with a flat shape. Therefore, although the range of eye movement within the orbit between species are different, it is difficult for most birds to move them freely (Fernández-Juricic, 2012). This limitation in eye movement leads to the result that birds primarily use their agile necks the move the head, rather than the eyes, to direct their gaze to the object they wish to pay attention to. Note that when looking at an object, birds orient the side faces to it instead of the frontal face (Tucker, 2000). Since the head direction approximately aligns with the gaze direction, to examine the perception of gaze, it is necessary to first investigate the perception of head direction.

In this research, we use Bengalese finches (*Lonchura striata* var. *domestica*) as the subject because they are monogamous songbirds, in which the parents take care of their offspring, and they also have close relationships with individuals outside their family, resulting in complex social behavior. Therefore, songbird is a suitable model animal for studying the perception of the gaze of other conspecific individuals.

We here confirm earlier findings on how songbirds recognize the head direction of other conspecifics. Before conducting behavioral experiments, we testified that the subjects could recognize the image stimuli presented on the monitor as the same species, which shows that our experiment setting is proper for this study. Hence, by presenting various directions between frontal and lateral images, we examined whether the subjects are able to discriminate between frontal and lateral images of conspecifics and whether there exists a categorical boundary.

## Materials and methods

### Animals

The subjects for this study were 9 female Bengalese finches (*Lonchura striata* var *domestica*). Birds were either purchased from a pet supplier or reared in our laboratory colony, and all were older than 10 months of age. Daily training and test sessions were done during the daytime (from 11 am to 2 pm). Food access was limited to 2–3 h around 3–6 pm to keep at 85% of their free-feeding weights, but vitamin-enriched water and shell grit were available ad libitum. The light/dark cycle was controlled to be constantly 14/10 h. Room temperature and humidity were maintained at approximately 25°C and 60%, respectively. The original research reported herein was performed under guidelines established by the Institutional Animal Care and Use Committee at the University of Tokyo.

### Apparatus

A test cage (W15.5 × D30.0 × H22.0cm) was placed in a soundproof box (inside dimension: W40 × D58 × H38cm) with LED lights inside on the top. The front panel had two response keys made up of transparent acrylic panels. One round key (diameter=10mm) served as the observation key, and the other square key (W5.5 × H4.0cm) served as a report key. The stimuli were presented on a liquid crystal display (IODATA, LCD-AD172SEB) located 1.5 cm behind the front panel, visible through the report key. The observation key was illuminated with a green color when active for pecking response. A feeder was placed on the cage and used to deliver grains into a food container located 5cm below the report key. A small light was used to illuminate the container for 2s when food had been delivered. We used a laptop computer (Dell, Latitude 3301) to operate the experiment and a web camera (Logitech, C505 HD720P) to record the birds’ behavior.

### Stimuli

The stimuli were shot from 6 adult Bengalese finches (3 males and 3 females, BF1-6) in an acrylic cage (W29.5 × D25 × H31.5cm) placed in a soundproof box (inside dimension: W57 × D48 × H48.5cm). We fixed the position of the bar lights on top of the cage to control the light environment of shooting. Each individual was videoed for 15-30 minutes with the 4K recording (iPhone XR) so that pictures of various face angles could be captured from the videos. Then, we imported pictures into a laptop computer (MacBook Pro Early2015) and cut them into the same size using Photoshop (Adobe).

The discrimination stimuli were 5 sets of frontal and right-side oriented images of stimuli birds (BF1-4, 6). The criteria for selecting the frontal images were symmetry and that both eyes be at approximately the same horizon. For the right-side images, the beak length was maximum in the entire video taken, and the direction of the beak was parallel to the perch in the upward webcam image. Before the test, only the head parts were cropped, and the backgrounds were changed to gray so that the background and other body parts in the image would not provide clues for discrimination.

Several images were cropped from the video to select three different face orientations between the frontal and right-side images for test stimuli. The criteria for filtering test stimuli from these alternative images were as follows. As shown in Figure 2, first, in each alternative image, the distance from the tip of the beak to the vertical line passing through the highest point of the beak was A. Thus, A’s value was the largest in the right-side images; the maximum value was recorded as b. The value of b differed depending on the individual. The A/b value of each alternative image was then calculated, and the image with those values of 0.2, 0.5, and 0.8 was selected as the test stimuli. The A/b values for the frontal and lateral images were 0 and 1.0, respectively. A left-right reversal of three test stimuli and right-side stimuli (discrimination stimuli) were added to the test stimuli. The A/b values for the left-oriented test stimuli were -0.2, -0.5, -0.8, and -1.0. The A and A/b values (px) and luminance data for each stimulus are summarized in Table1.

**Figure 1.**
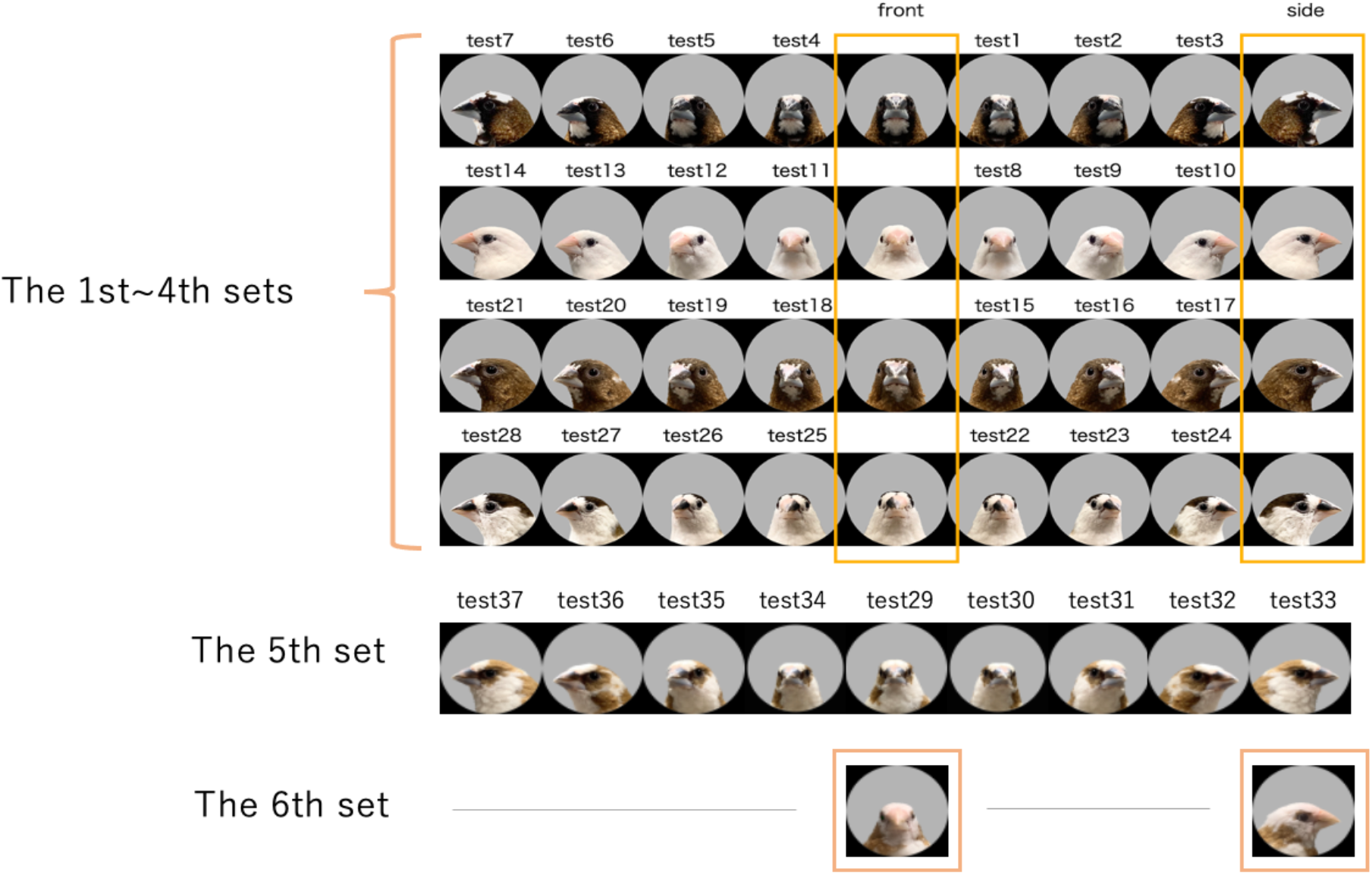
All stimuli of Bengalese finches’ faces in various orientations. Images in the orange frame were the discrimination stimuli, and the others were test stimuli.

**Figure 2.**
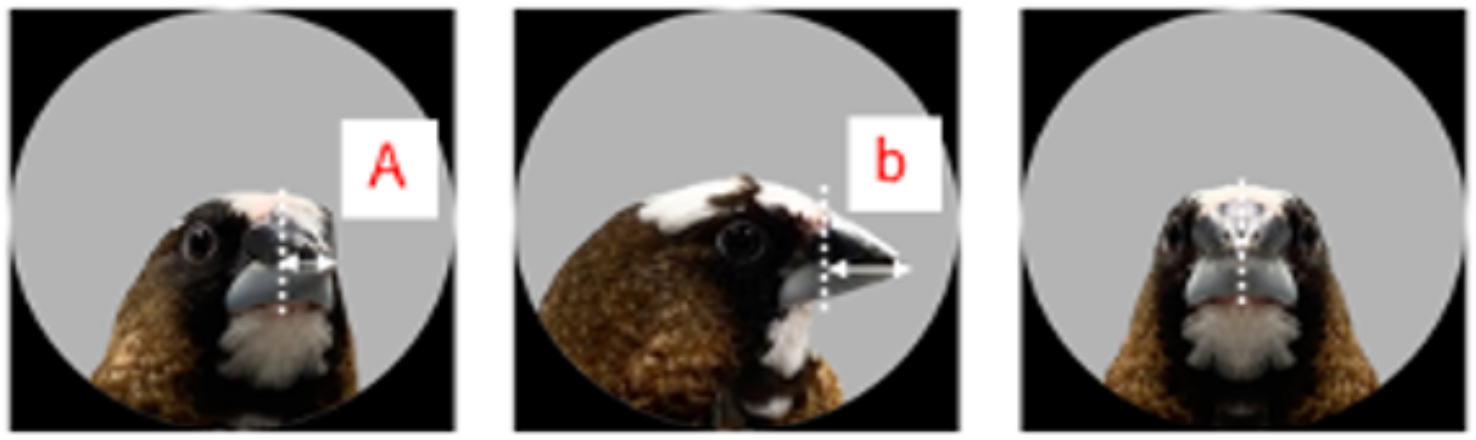
The selection criteria of test stimuli.

**Figure 3.**
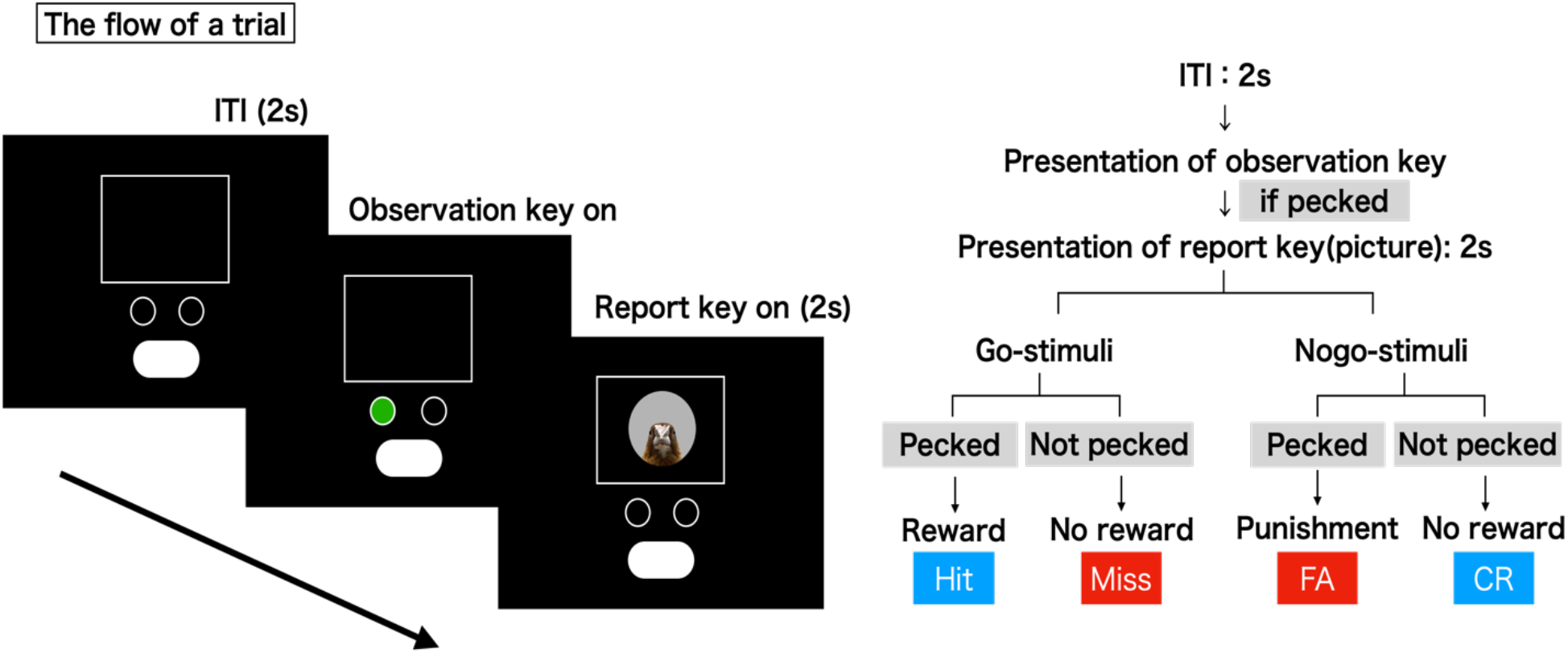
GO/NOGO task flow of one trial.

**Figure 4.**
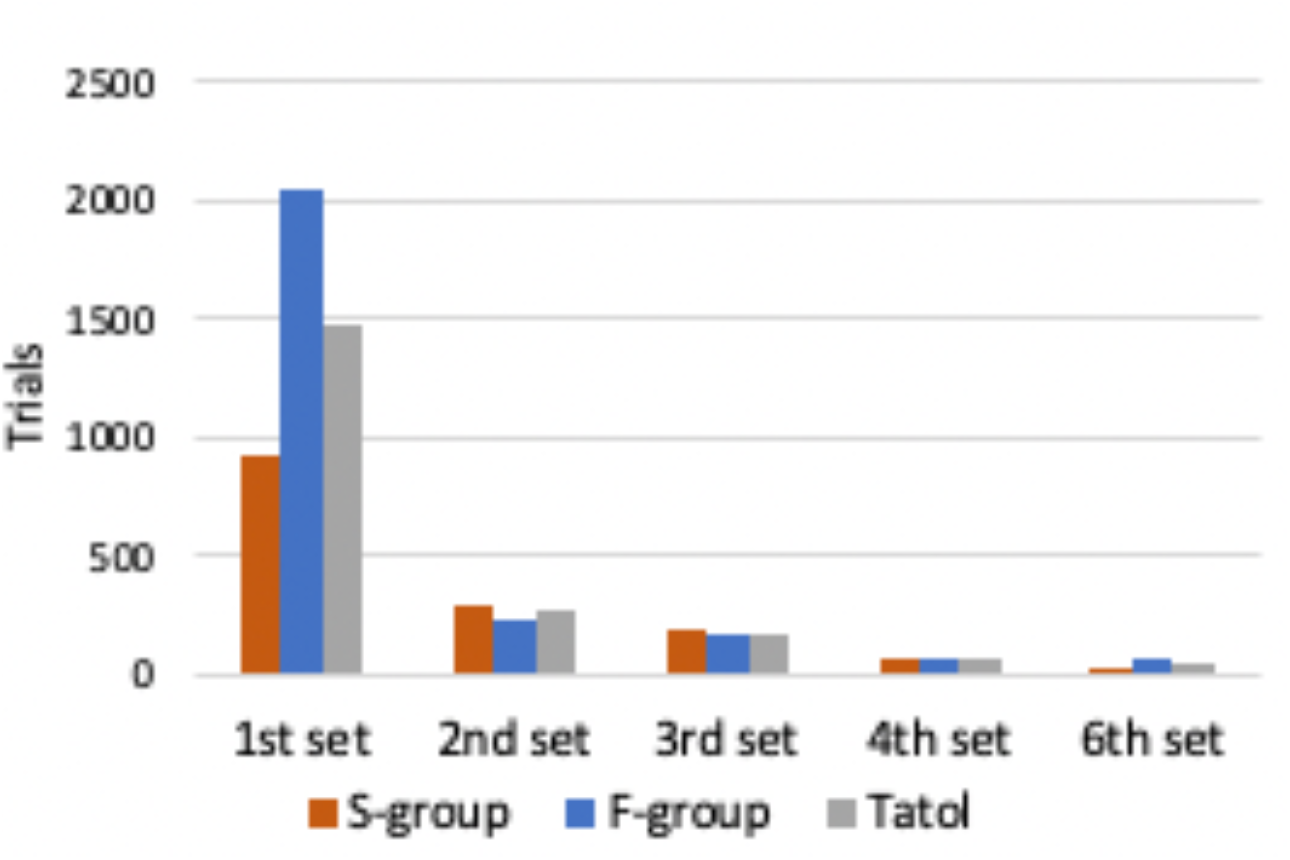
Numbers of trials until the novel stimuli set can be discriminated.

### Procedure

#### verification experiment

To confirm whether the subjects recognized the images presented by the apparatus as the same species, we showed a slide of the same and other animal species for approximately 5 minutes and observed birds’ behaviors. Two birds (one male and one female) were used in the verification experiment.

#### Operant conditioning

First, subjects were trained to peck the observation key using an auto-shaping method. After the subjects learned to peck the observation key spontaneously, they were trained to peck the report key within 2s after triggering the observation key. Then a GO/NOGO task was applied, in which we trained the subjects to discriminate between frontal and right-side oriented images. All subjects were divided into two groups, one with frontal images as the GO stimuli(F-group) and the other with side images as the GO stimuli(S-group).

The observation key was activated as a signal at the beginning of each trial. A GO or NOGO stimulus was presented in a pseudorandom order following an observation key peck. Next, the report key was activated, and the stimulus was presented simultaneously. After a GO stimulus, pecking the report key within 2s resulted in a food delivery (Hit); otherwise, the bird did not obtain a reward for the trial (Miss). After a NOGO stimulus, pecking the report key within 2s resulted in a punishment of blackout for the 20s (False Alarm; FA); otherwise, the bird proceeded to the subsequent trial without any consequence (Correct Rejection; CR). The inter-trial interval was 2s.

At first, subjects were required to discriminate against one pair of discrimination stimuli. When the subject is correct at 90 percent, we add one more pair of stimuli until up to four. If there is no significate difference in the bird’s performance for three consecutive training sessions, correction trials were applied for unsuccessful trials (FA or Miss) until the bird responded correctly. The training was conducted 5-6 days a week, one session per day, with 144 trials (72 each for GO and NOGO trials). Before the experiment was suspended due to the spread of covid-19, 100 trials per session were used. Each training session concluded when all trials were completed or 1 hour after starting the session.

The test session consisted of 63 trials each of GO and NOGO stimuli used in training (The first 24 trials at the beginning of a session were warming-up trials), and 18 trials presented test stimuli, for a total of 144 trials. Both training and test trials were presented in a pseudorandom order such that the same stimulus was not presented more than three times in a row. Each test stimulus was presented 20 times. If the insertion of test stimuli caused performance on a training trial to drop below 80%, the subject was retrained until the learning criterion of 90% correct response rate was exceeded.

## Results

We found that the male Bengalese finches sang to the individual presented in a video by a liquid crystal display. At the same time, females approached the side where the video stimuli were given or behaved actively on the perch. In addition to the stimuli of Bengalese finches, we also presented a slide show with images of a Java sparrow, hawk, dog, and cow that changed every 15 seconds. As a result, we observed the animals flapping their wings violently and clinging to the fence in the experimental cage due to their fear of the images of other species

For the discrimination task, 12 trials were counted as one block, and the correct response rate (CR+Hit/CR+Hit+FA+Miss) was calculated for each block. 9 out of 8 birds could discriminate all stimuli. One bird was excluded from the analysis and test task because it could not discriminate the first pair of stimuli after three months of training. The mean trials number of the group that used the frontal images as the GO stimuli (F-group), the group that used the side images as the GO stimuli (S-group), and all subjects are shown in Figure 5. The results showed a significant difference in the number of trials required as the number of pairs discriminated increased (one-way ANOVA, F = 11.027, p <.05). Comparing the mean number of trials needed by the two groups, it was found that the F-group required more trials than the S-group, especially in discriminating the 1^st^ set of stimuli (mean number of trials for the F-group: 2042, 234, 174, 57 trials; mean number of trials for the S-group: 921, 300, 180, 66 trials).

**Figure 5.**
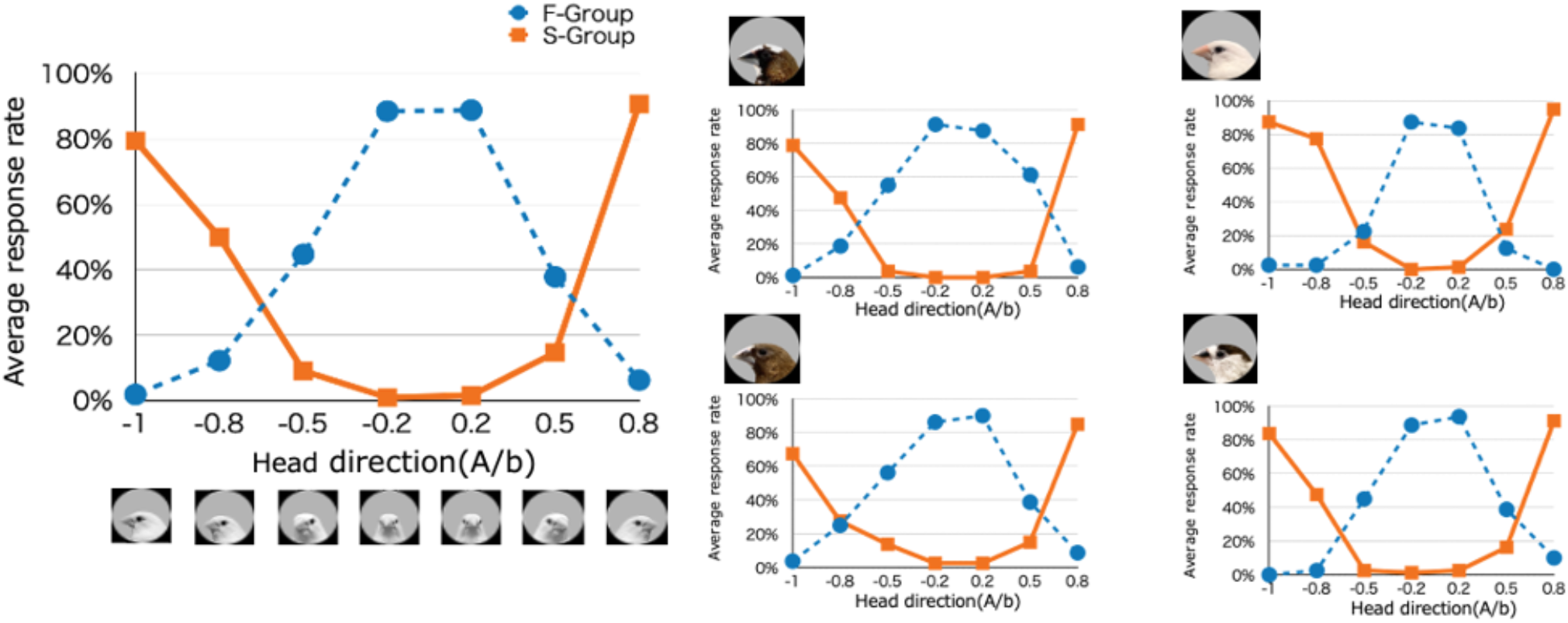
Average response in the test (1st-4th sets).

**Figure 6.**
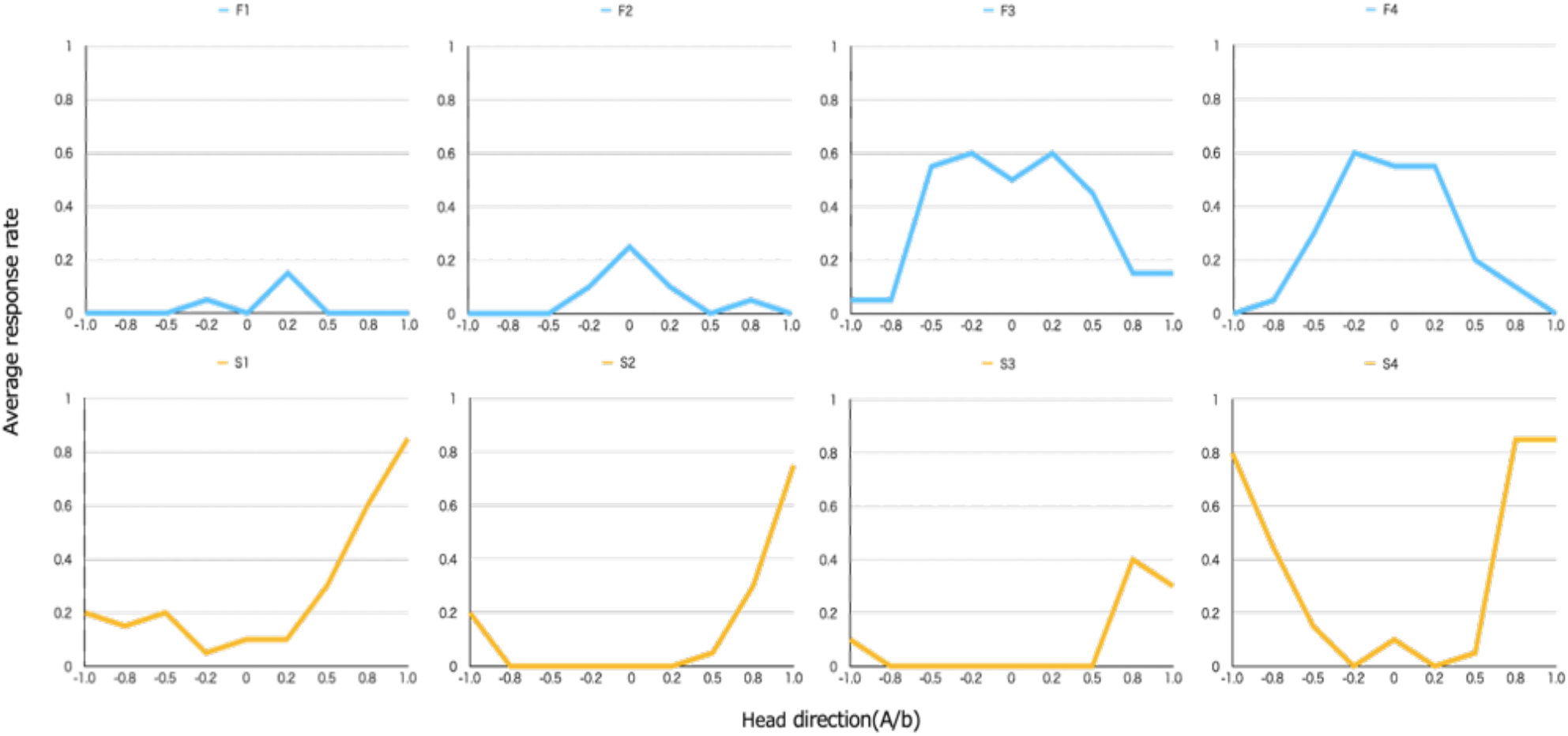
The average response of the 5th sets by each subject(F-group: F1-4, S-group: S1-4).

The average response rate was plotted for the test as the number of times subjects were pecked to each test stimulus by the total number of presentations (4 subjects x 20 times). The average response rate to each test stimulus for all individuals is shown in Figure 7. Some response rates were categorical, such as the 2^nd^ set, but others were continuous, such as the 1^st^, 3^rd^, and 4^th^ pairs. (A sharp drop in response rate from 77.5% to 16.3% when the A/b value changed from -0.8 to -0.5, and a sharp increase in response rate from 23.8% to 95.0% when the A/b value changed from 0.5 to 0.8.) The average response rate to the four sets of stimuli showed that the Bengalese finches had a relatively continuous response to the seven head orientations.

## Discussion

We observed that male Bengalese finches emitted directed song as a sexual display towards the images presented by the liquid crystal display, which confirmed the results reported by Ikebuchi&Okanoya (1999). Based on the above observations, we assumed that the animals recognized the images of the Bengalese finch presented by the monitor in the experimental apparatus as a living conspecific individual.

The result of the discrimination task indicates that as the number of sets increased, the subjects needed fewer trials to achieve the criterion. This suggests birds could use the information they had already acquired to quickly complete the novel discrimination task. Our experiment shares a similar procedure with the learning set experiment. The most common procedure for explicitly testing the formation of learning sets is to give subjects several different discrimination tasks successively and measure how the number of incorrect responses decreases as the number of tasks increases (Bitterman, 1975). In the learning set paradigm, animals are trained on a series of simple discrimination tasks containing a pair of stimuli, where one is delivered with reinforcement and the other is not. In this research, for example, the subjects in F-group are reinforced if they choose the frontal image, and no reinforcement is given if they choose the lateral image. In general, the resolution of the initial discrimination in the learning set paradigm is accomplished through trial and error, but the subject’s behavior appears insightful.

After solving many problems similar to the initial discrimination, skilled subjects performing such discrimination tasks behave as following a win-stay/lose-shift strategy (Nowak & Sigmund, 1993). That is, (1) on the first trial anyway, they randomly choose one of the stimuli and check the outcome. Food may or may not be given according to whether the choice is correct or incorrect. (2) If the choice were correct, the subjects would choose that stimulus again the next time, and if the choice were incorrect, they would shift to another stimulus (Harlow, 1949). In our GO/NOGO task, (1) the subjects pecked at all stimuli, regardless of the stimuli type, when discriminating the 1^st^ set of discrimination stimuli. Pecking at the GO stimulus was reinforced with food, while pecking at the NOGO stimulus was punished with a 20s blackout. (2) Subjects learned not to peck either of the discrimination stimuli because they were immediately moved to the subsequent trial if they did not peck the NOGO stimulus within 2s. The same trial-and-error process was repeated for the 2^nd^-4^th^ sets of discrimination stimuli. The birds responded differently to the frontal and lateral images of the novel stimuli set. As they became more proficient, especially when the 4^th^ set was added, they required only a minimal number of trials to achieve the learning criterion. Furthermore, it is possible that when moving to a new task, the subjects gained insight into the relationship corresponding to the preceding stimuli set, which may have contributed to the reduction in the number of incorrect responses. They tended to peck at one stimulus with similar characteristics to the 1^st^ GO stimuli and not to peck at the other stimuli that took similar attributes to the NOGO stimuli. Specifically, if the first set of GO stimuli were a frontal image, the participants would peck at the same frontal image when presented with the novel stimuli set, even though the feather color differed. As the discrimination task progressed, it is possible that all discrimination stimuli were divided into two categories. More novel stimuli set need to be inserted to verify the establishment of the two categories of frontal and lateral.

In the test, except for discrimination stimuli, we examined responses to seven different head orientations of test stimuli (three right-facing and four left-facing lateral images). Ignoring individual differences among stimuli subjects, we found that the average response rate to these test stimuli was unexpectedly continuous. There may be no clear discrimination boundary when birds discriminate against the head orientations of other conspecific individuals. Why those which were especially good at dividing frontal and lateral faces responded non-categorically towards various direction is less clear. First, subjects may not perceive others’ face orientation in a categorical way claimed by Liberman et al. (1957), or only perform category perception toward specific individual such as court mate and kinship. If the phenomenon of “speech categories” or “phoneme categories” did not exist in gaze recognition, it would be natural that the subjects’ perception of head orientations would be continuous, with no sudden reversal in their responses after a particular direction. In operant conditioning, when a specific response occurs to a specific stimulus, the same response will occur to similar stimuli, which is called generalization (Davydov, 1990). Conversely, when the orientations are identical to that of the NOGO stimulus, no pecking occurs, the mean response rate drops, and an overall continuous curve is obtained. It is difficult to infer from the results of this experiment alone that subjects perceive the head direction of other individuals categorically. It is necessary to show that up to a certain level of variation, birds perceive the area occupied by their beaks and the position of their eyes on their heads as frontal or lateral, even though the stimuli are physically slightly different. The effect of category perception on discrimination and discrimination judgments in humans, in general, is called category perception. One way category perception influences discrimination is that differences are not perceived as readily when comparing image stimuli within the same frontal or lateral category but are perceived more clearly when comparing image stimuli within different categories.

The second factor could be the inaccuracy of quantifying the image stimuli concerning head orientations or the small number of head orientations used for the test stimuli, which resulted in only incomplete curves that appeared to be continuous. Plotting the average response rate for each test stimuli revealed a point at which the average response rate changed abruptly at two adjacent orientations. However, because of the small number of different test stimulus orientations in this study, it is not possible to determine whether that was the orientation at which the response changed abruptly or whether the continuous change appeared to be locally abrupt. In the four test stimulus sets classified as the same orientation, the error between the stimuli needs to be further reduced.

The response rates to the seven test stimuli extracted from physically continuous head orientations between frontal and lateral views yielded continuous curves. However, the range of frontal or lateral recognition is not uniform, and it is not possible to determine whether there is a boundary line for discrimination or not. In future work, we need to increase the number of test stimuli to clarify further the categorical perception of the head orientations of other individuals.

Comparing the average number of trials required to achieve the learning criterion for both groups in the discrimination task showed that the F-group had more trials than the S-group, especially in the 1^st^ pair of discrimination stimuli. The difference in results between the two groups indirectly supports that the Bengalese finches perceive stimuli as homologous rather than neutral images. The differences between the two groups indirectly support that the Bengalese finches perceive the image stimuli as the same species rather than neutral images. The learning set was not as pronounced for the second and subsequent sets of image stimuli. After the learning set is formed, these ecological factors have less influence on discrimination performance.

Birds efficiently learned how to discriminate novel stimuli once they were able to discriminate stimuli set through trial and error may provide evidence supporting the formation of learning sets. This problem-solving ability allows animals to use their previous experience to obtain more rewards at less cost when faced with a new situation. Riddell & Corl (1977) published learning set performance in various mammals, including primates and carnivorous rodents. The data showed that learning set performance was highly correlated with Nc, a measure of the number of cortical neurons in mammals that estimates the larger-than-necessary portion of their size. Although animals differ in their sensory-perceptual abilities, this correlation is indicative of the relationship between the mammalian neocortex and the ability of the neocortex to process visual information. However, since it indicates that birds, which do not possess a neocortex, may also have higher-order information-processing abilities, it may be that the prototypical neurons of the neocortex were already present in the common ancestral stages of mammals and birds or that these abilities were acquired during their respective evolutionary paths.

From the test task, when the physical quantity is presented as a continuously varying head orientations, the birds represent a continuous average response rate to the image stimulus. Considering the head orientations as an approximation to the gaze directions, it is highly possible that the head orientations are not uniformly classified into two categories for the gaze of other individuals. Then, in judging the attention of other individuals, they may be perceiving it in combination with sound signals and other signals commonly used in animal communication. In addition, the subjects used for image stimuli in this experiment were strangers, not living together. Therefore, it is expected that using an individual kept in the same cage as the subject would increase the motivation for discrimination and make the perception of head orientations more bipolar.

## Acknowledgements

This work was supported by MEXT/JSPS Grant-in-Aid for Scientific Research on Innovative Areas #4903 (Evolinguistics), Grant Number JP17H06380.

## Notes

### Competing Interest Statement

The authors have declared no competing interest.

## References

1. Bitterman, M. E. (1975). The comparative analysis of learning. Science, 188(4189), 699–709.

2. Davydov, V. V. (1990). Types of Generalization in Instruction: Logical and Psychological Problems in the Structuring of School Curricula. Soviet Studies in Mathematics Education. Volume 2. National Council of Teachers of Mathematics, 1906 Association Dr., Reston, VA 22091.

3. Dukas, R. (2002). Behavioural and ecological consequences of limited attention. Philosophical Transactions of the Royal Society of London. Series B: Biological Sciences, 357(1427), 1539–1547.

4. Fernández-Juricic, E. (2012). Sensory basis of vigilance behavior in birds: synthesis and future prospects. Behavioural Processes, 89(2), 143–152.

5. Ikebuchi, M., & Okanoya, K. (1999). Male zebra finches and Bengalese finches emit directed songs to the video images of conspecific females projected onto a TFT display. Zoological Science, 16(1), 63–70.

6. Haque, A., & Dickman, J. D. (2005). Vestibular gaze stabilization: different behavioral strategies for arboreal and terrestrial avians. Journal of Neurophysiology, 93, 1165–1173.

7. Harlow, H. F. (1949). The formation of learning sets. Psychological review, 56(1), 51.

8. Hughes, A. (1977). The topography of vision in mammals of contrasting life style: comparative optics and retinal organisation. In The visual system in vertebrates (pp. 613–756). Springer, Berlin, Heidelberg.

9. Liberman, A. M., Harris, K. S., Hoffman, H. S., and Griffith, B. C., Journal of Experimental Psychology, 54 (5), 358–368, 1957.

10. Nowak, M., & Sigmund, K. (1993). A strategy of win-stay, lose-shift that outperforms tit-for-tat in the Prisoner’s Dilemma game. Nature, 364(6432), 56–58.

11. Pratt, D. W. (1982). Saccadic eye movements are coordinated with head movements in walking chickens. Journal of Experimental Biology, 97, 217–223.

12. Santos, L. R., & Hauser, M. D. (1999). How monkeys see the eyes: Cotton-top tamarins’ reaction to changes in visual attention and action. Animal Cognition, 2(3), 131–139.

13. Tomasello, M., & Carpenter, M. (2007). Shared intentionality. Developmental science, 10(1), 121–125.

14. Tucker, V. A. (2000). The deep fovea, sideways vision and spiral flight paths in raptors. Journal of Experimental Biology, 203(24), 3745–3754.

15. Walls, G. L. (1942). The vertebrate eye and its adaptive radiation. Bloomfield Hills, MI: Cranbrook Press.

16. Wohlschläger, A., Jäger, R., & Delius, J. D. (1993). Head and eye movements in unrestrained pigeons (Columba livia). Journal of Comparative Psychology, 107, 313.

17. Yorzinski, J. L., Patricelli, G. L., Babcock, J. S., Pearson, J. M., & Platt, M. L. (2013). Through their eyes: selective attention in peahens during courtship. Journal of Experimental Biology, 216(16), 3035–3046.

18. Yorzinski, J. L., Patricelli, G. L., Platt, M. L., & Land, M. F. (2015). Eye and head movements shape gaze shifts in Indian peafowl. Journal of Experimental Biology, 218(23), 3771–3776.

